# Rapidly decreasing phasic dopamine responses precede a hedonic switch during satiation of sodium appetite

**DOI:** 10.64898/2026.07.16.738888

**Authors:** Paula Bazzino, Dionne CR Russchen, Alexandra T Keinath, Mitchell F Roitman, James E McCutcheon

## Abstract

**Highlights:**

- Phasic dopamine tracks changing physiological need during sodium appetite
- Rapid decreases in dopamine precede the loss of appetitive behavioural responses
- Transitions occur abruptly within individuals despite gradual group averages
- Dopamine responses track cumulative sodium intake during sodium satiation

Phasic dopamine signalling updates the value of stimuli and actions. Most empirical support for this critical process involves manipulation of extrinsic stimuli (e.g., reward magnitude, reward omission). Yet interoceptive signals clearly influence motivated behaviour. Sodium depletion generates a sodium appetite where the value of sodium increases. How the value of sodium is updated during ingestion and as animals satiate their need remains unknown. Here, we administered sodium chloride solutions via intraoral delivery and measured appetitive behaviour and dopamine release in sodium replete and deplete rats. Two different concentrations of sodium chloride were used to modulate the rate of repletion for sodium deplete rats. Appetitive behaviour was evoked by infusions only when rats were deplete, and this behavioural motif faded away during the session only when high concentration infusions were delivered. Phasic dopamine release in the nucleus accumbens had a similar pattern. We identified transition points for both behaviour and dopamine in the subgroup of rats that reached satiation and found that transitions were sharp, with dopamine decreasing just before behaviour. Moreover, transitions occurred only after enough sodium was infused to replace that typically lost upon depletion. These findings demonstrate that mesolimbic dopamine dynamically updates the value of taste stimuli based on interoceptive signals that relay satiation.

## Introduction

Sensing physiological need (e.g., calories, fluid) is essential for counterregulatory autonomic responses and to motivate behaviour. But knowing when to say ‘when’ - that is, sensing satiation - is equally critical, as overconsumption can threaten health. A striking example is the crucial need to maintain plasma sodium within a tight range (Geerling & Loewy, 2008; Grove & Knight, 2024). Sodium loss induces an innate and specific sodium appetite (Rowland, 2023), which is highly-conserved across phylogenetically diverse species, from insects to humans (Beauchamp et al., 1990; McDowell et al., 2026). Overconsumption of sodium, though, can lead to hypertension and other life-threatening complications (Farquhar et al., 2015). The induction of sodium need, by deprivation or depletion, causes a hedonic switch (Berridge et al., 1984) and induces seeking and appetitive behaviour directed at the very same high concentration (i.e., hypertonic) sodium solutions that would otherwise be avoided or rejected (Breslin et al., 1993; Quartermain et al., 1967; Robinson & Berridge, 2013). Thus, sodium appetite represents a taste by state interaction where the value of sodium increases only when it is needed.

Across multiple homeostatic needs (e.g., hunger, thirst, sodium appetite), the peripheral factors and central circuits responding to deficit are well understood (Lowell, 2019). Included in this process is the recruitment of the mesolimbic dopamine system (Roitman & McCutcheon, 2025). Brief, high concentration (phasic) dopamine release in the nucleus accumbens is evoked by the taste of hypertonic sodium but only when animals are sodium deplete (Cone et al., 2016; Fortin & Roitman, 2018). In turn, phasic dopamine activity is thought to be critical in assigning value to stimuli (Bornhoft et al., 2025) and actions (Bakhurin et al., 2025; Coddington et al., 2023). Homeostatic reinforcement theory suggests that dopamine and the value of sodium are updated by decreasing drive during ingestion (Duriez et al., 2023). However, this has not yet been empirically demonstrated. Indeed, little is known about how ongoing ingestion and repletion impact phasic dopamine signalling and ultimately behaviour.

There are two critical barriers that hinder the study of dopamine in devaluation due to satiation. First, for most stimuli, valence does not depend on state, e.g., sugar remains positive even with repletion. Second, animals reduce approach and consumption with satiation making it difficult to dissociate value from action. Hypertonic sodium ingestion in the sodium deplete rat offers a unique opportunity to overcome these challenges. Here, we measured dopamine release and behavioural responses in rats repeatedly tasting saline as their sodium need state varied. Specifically, we assayed dopamine release in the lateral shell of the nucleus accumbens via GRAB-DA fibre photometry while tracking the expression of appetitive behaviour. To ensure that we could continue to assess both dopamine and behaviour even as rats became satiated, saline was delivered directly into the mouth via an intraoral catheter. We manipulated homeostatic need prior to each session (deplete versus replete), and varied the rate of repletion by using saline of different sodium concentrations (100 mM versus 450 mM). Our data demonstrate that abrupt within-session decreases in dopamine release are a function of the amount of sodium delivered and actually precede changes in behaviour by just a few trials, supporting the transition from appetitive behaviour when in need to aversive behaviour in satiation.

## Results

### Assaying sodium-evoked neural and behavioural responses as physiological need state varies

One challenge in assaying neural and behavioural responses to sodium across physiological states is that animals avoid sodium when needs have already been met. To overcome this limitation, we implanted each rat (*n* = 20; female = 10) with an intraoral catheter allowing delivery of NaCl directly to the oral cavity (Figure 1A). To assay dopamine release evoked by intraoral infusion of NaCl, we expressed GRAB-DA1h in the lateral shell of the NAc and implanted a fibre to gain optical access (Figure 2A). To characterise behaviour, we simultaneously recorded a birds-eye video of the rat during all experiments.

**Figure 1.**
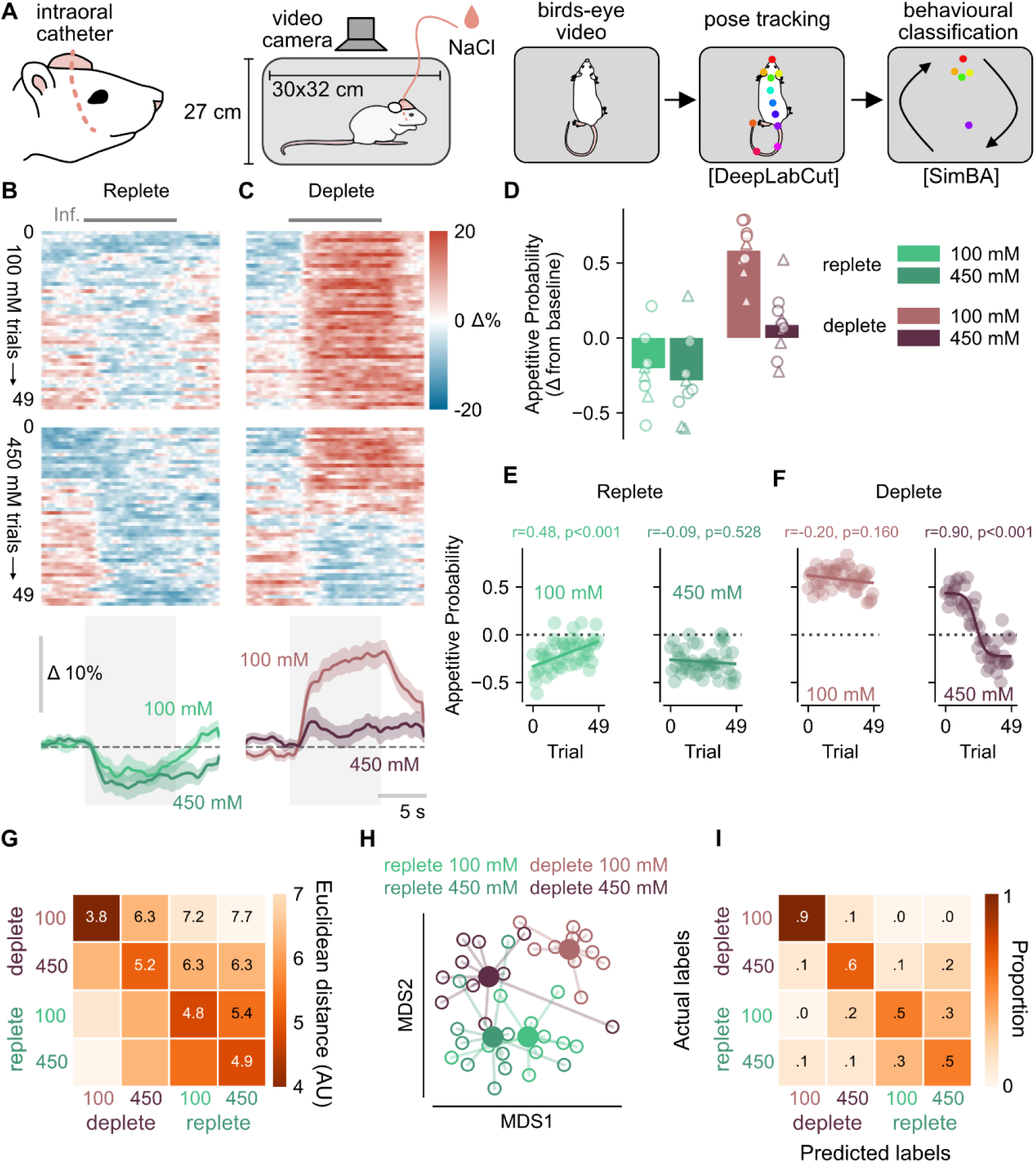
Behavioural responses to NaCI are dependent on physiological state and NaCI concentration. (A) Schematic showing experimental setup for behavioural measurements. Rats were given intraoral NaCI while being videoed from above. DeepLabCut was used to extract body parts and SimBA was used to detect appetitive behaviour. (B) Heatmaps showing probability of appetitive behaviour in sodium replete rats given 10-second intraoral infusions of low (100 mM; upper heatmap) or high (450 mM; lower heatmap) concentration of NaCI. Time of infusion indicated by gray bar above heatmap. Bottom panel shows mean ± SEM. n=10 rats in each group. (C) As in (B) but in the same rats after sodium depletion. (D) Mean behavioural response averaged across all trials in session. Bars are mean and markers are individual rats (circles, male; triangles, female). (E) Mean behavioural response per trial in sodium replete rats. Circles show mean from n=10 rats and line shows linear fit to data with statistics shown above plot (F) As in (E) but for rats when sodium depleted. For 450 mM rats, data were best fit with a sigmoidal function. (G) Heatmap showing mean similarity (Euclidean distance) between behavioural responses of different groups with darker colours representing closer euclidean distances (H) Multidimesional scaling (MDS) used to show distances between behaviour of individual rats (open circles) and mean of each group (filled circles). (I) Confusion matrix showing classification accuracy for behavioural responses for different groups where rows are true class and columns are predicted class.

**Figure 2.**
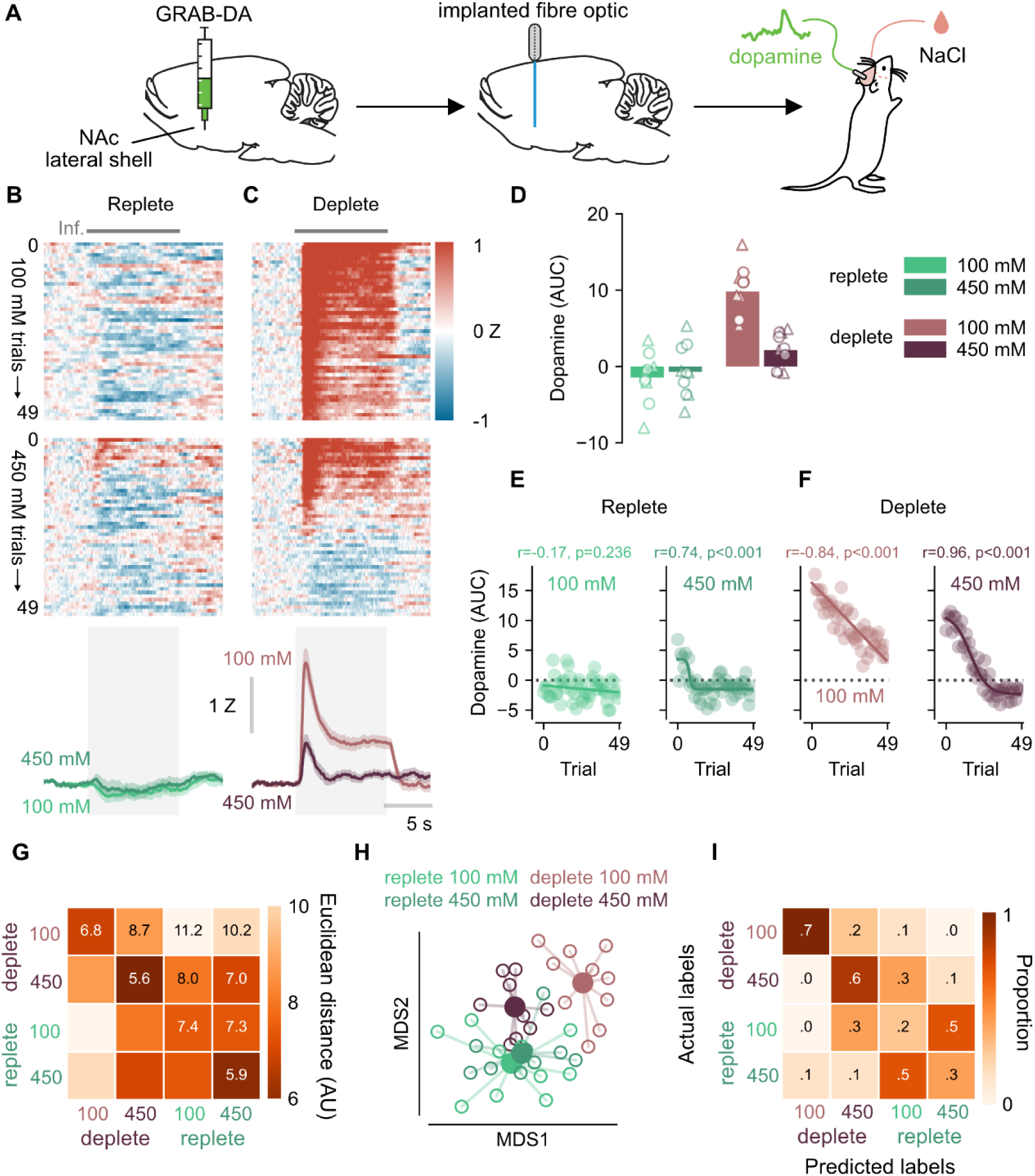
Dopamine responses to NaCI are dependent on physiological state and NaCI concentration. (A) Schematic showing methods for fibre photometry of dopamine release. GRAB-DA was expressed in nucleus accumbens lateral shell and measured via implanted fibre optic during NaCI infusions. (B) Heatmaps showing dopamine release in sodium replete rats given 10 s intraoral infusions of low (100 mM; upper heatmap) or high (450 mM; lower heatmap) concentration NaCI. Time of infusion indicated by gray bar above heatmap. Bottom panel shows mean ± SEM. n=10 rats in each group. (C) As in (A) but in the same rats after sodium depletion. (D) Mean AUC of dopamine response averaged across all trials in session. Bars are mean and markers are individual rats (circles, male; triangles, female). (E) Mean AUC per trial in sodium replete rats. Circles show mean from n=10 rats and line shows linear fit to data with statistics shown above plot. (F) As in (E) but for rats when sodium depleted. For 450 mM rats, data were best fit with a sigmoidal function. (G) Heatmap showing mean similarity (euclidean distance) between dopamine responses of different groups with darker colours representing closer euclidean distances. (H) Multidimesional scaling (MDS) used to show distances between dopamine responses of individual rats (open circles) and mean of each group (filled circles). Confusion matrix showing classification accuracy for dopamine responses for different groups where rows are true class and columns are predicted class

During each experimental session, rats received 50 intraoral infusions (‘trials’) of NaCl, each lasting ten seconds and separated by variable intertrial intervals. Rats could choose to ingest or reject each infusion, but were nevertheless required to taste the solution. The volume of each infusion was fixed (400 µL) but the NaCl concentration was varied between two groups of rats: One group received a low concentration of NaCl (hypotonic; 100 mM) and one received a high concentration (hypertonic; 450 mM). Each rat was tested with its designated concentration in both sodium replete (R) and sodium deplete (D) conditions. Sodium depletion was accomplished through the administration of the diuretic furosemide 1 day prior to recording (see Methods for further details). For convenience we refer to each of these groups according to their combination of condition and concentration by abbreviations (i.e., Replete, 100 mM infusions: **R100**; Replete, 450 mM infusions: **R450**; Deplete, 100 mM infusions: **D100**; Deplete, 450 mM infusions: **D450**).

### Sodium-evoked behavioural expression reflects physiological need

To characterise behavioural reactivity with high precision as physiological need varied, we employed a multistep machine learning pipeline. First, DeepLabCut was trained to track multiple body parts across each session. As an initial analysis, we focused purely on the movement of the head as traditional taste reactivity procedures use head shakes as a way of identifying responses to aversive stimuli (Berridge et al., 1984; Grill & Norgren, 1978). Plotting infusion-evoked head movement across trials (Figure S1) revealed clear group differences across combinations of physiological condition and infusion concentration. Replete rats (R100 and R450) exhibited small increases in head movement during infusions, while deplete rats (D100 and D450) exhibited decreases in head movement. For D100 rats, infusion-evoked decreases in head movement persisted throughout the session, while for D450 rats this behavioural motif weakened across trials.

To better discriminate between infusion-evoked behaviours, we next trained a classifier to recognise appetitive-like behaviour from the DeepLabCut-derived poses (SimBA; Goodwin et al., 2024). Briefly, the classifier was trained using D100 rats based on the assumption that the infusions remained appetitive across all trials for these rats, consistent with the prior head movement analysis. After training this model via leave-one-out cross validation (see Methods), we applied it to quantify the probability of appetitive behaviour throughout each session for all rats.

This analysis yielded qualitatively similar group differences as observed with head movement analysis, while providing a more robust characterisation of appetitive responding (Figure 1 and Figure S2). D100 rats exhibited an increased probability of appetitive behavioural expression during infusions whereas both R100 and R450 rats exhibited a decrease in appetitive behaviour during infusions. Notably, D450 rats appeared to change their behavioural expression as the session progressed, with most showing a high infusion-evoked probability of appetitive behaviour at the start of the session and a low probability by the end (Figure 1C). Averaging responses across trials showed that infusions of either sodium concentration decreased the probability of appetitive behaviour in replete rats (Wilcoxon signed-rank test; R100: *W* = 6, *p* = 0.027, *n* = 10; R450: *W* = 4, *p* = 0.014, *n* = 10; Figure 1D), whereas infusions increased appetitive behaviour in D100 rats (*W* = 15, *p* = 0.002, *n* = 10). The averaged response for D450 rats did not differ from zero (*W* = 0, *p* = 0.232, *n* = 10) likely due to the change in behaviour across trials during the session. A mixed effects ANOVA revealed a main effect of Condition (*F*_1,16_ *=* 162.67, *p* < .001), Concentration (*F*_1,16_ = 11.04, p = 0.004) and a Condition x Concentration interaction (*F*_1,16_ = 21.10, *p* < .001). There was no main effect of Sex or any significant interactions with Sex (*p*s > 0.05). Pairwise Holm-corrected post hoc tests showed that all groups differed from each other (*p*s < 0.05) except for the two replete groups (R100 vs R450, *p* = 0.483). This pattern of results demonstrates that sodium depletion recruits the expression of appetitive behaviour to the taste of sodium, including to a hypertonic solution.

Next, to better characterise trial by trial changes in appetitive behavioural expression, we plotted appetitive probability across trials for each group and examined the function which best described each (Figure 1E and 1F). R100, R450, and D100 rats were all best fit by linear regression (Table S1), reflecting minimal and/or gradual changes in appetitive expression across trials. On the other hand, the appetitive expression of D450 rats was best described by a sigmoid, reflecting an initial recruitment of appetitive behaviour to hypertonic saline followed by an abrupt decline in appetitive behaviour midway through the session (trial of transition midpoint (x_0_) from sigmoid fit = 24). Rats lose, on average, ∼2.5 to 3 milliequivalents of sodium following depletion with furosemide (Krause et al., 2010; Lundy et al., 2003). The transition point for D450 rats occurred after infusion of ∼4.3 milliequivalents of sodium, consistent with the change in appetitive behaviour following the development of post-ingestive satiating signals (Krause et al., 2010; Roitman et al., 1997). Together, these results demonstrate that the expression of appetitive behaviour is dynamic within session and is impacted by ongoing ingestion.

Finally, we examined whether sodium-evoked behavioural responses could reliably distinguish rats across physiological states. To do so, we first computed the euclidean distance between pairwise comparisons of rats based on the probability of appetitive behaviour across infusion periods (see Methods; Figure 1G and S2C). Next, we used multidimensional scaling to visualise the dissimilarity structure in two dimensions (Figure 1H). This analysis revealed that rats within each condition were generally more similar (i.e., closer) to one another than to rats in other conditions. R100 and R450 rats were similar to one another, while D100 rats differed greatly from all other groups. The cluster of D450 rats fell in between these two extremes. This pattern is consistent with the within-session behavioural transition seen in this group as sodium need was replenished throughout the session. Quantifying these observations, we found that a cross-validated classifier could identify the group to which rats belonged based on their sodium-evoked behaviour across trials with accuracy well above chance (accuracy = 62.5%; chance = 25.0%; binomial test *p* < 0.001; see Methods). Classifier confusions were most frequent between the replete groups, reflecting the poorer separation between replete R100 rats versus R450 rats (Figure 1I). Together, these results demonstrate that sodium-evoked behavioural expression reliably reflects physiological need, even as that need wanes with ingestion.

### Sodium-evoked phasic dopamine release reflects physiological need

To determine whether sodium-evoked phasic dopamine release varied across physiological states, we examined dopamine release in NAc lateral shell during infusions via fibre photometry (Figure 2A and S3). These data revealed a pattern of infusion-evoked phasic dopamine release which echoed the behavioural characterisation described above. In the replete state, dopamine responses were minimal to either concentration (Figure 2B). In contrast, in the deplete state, low concentration infusions evoked large dopamine responses throughout the session, whereas high concentration infusions elicited large dopamine responses in early trials, which diminished as the session progressed (Figure 2C).

To quantify these observations, we first computed the area under the curve (AUC) of the dopamine signal during the infusion period and averaged the AUC across all trials for each rat. This analysis revealed that, when rats were replete, neither concentration evoked reliable average dopamine release (Wilcoxon signed-rank tests versus zero: R100: *W* = 13, *p* = 0.160, *n* = 10; R450: *W* = 20, *p* = 0.492, *n* = 10). In contrast, D100 rats exhibited large average dopamine release (*W* = 0, *p* = 0.002, *n* = 10), while D450 rats exhibited a smaller but still significant average dopamine release (*W* = 6, *p* = 0.027, *n* = 10). Mixed effects ANOVA revealed a main effect of Condition (*F*_1,16_ = 73.23, *p* < 0.001), Concentration (*F*_1,16_ = 8.25, *p* = 0.011) and a significant Condition x Concentration interaction (*F*_1,16_ = 26.03, *p* < 0.001). There was no main effect of Sex or any interactions with Sex (*p*s > 0.05). Holm-corrected post hoc tests showed that dopamine responses in D100 rats differed from all other groups (*p*s < 0.001), dopamine responses in D450 rats trended towards differing from the replete groups (*p* = 0.074 vs. R100 and *p* = 0.074 vs. R450 rats), and R100 and R450 rats did not differ from each other (*p* = 0.636). This pattern of results demonstrates that sodium depletion recruits phasic dopamine release to the taste of sodium, including to a hypertonic solution.

To better characterise trial by trial changes in sodium-evoked phasic dopamine release, we plotted the AUC of the dopamine signal during the infusion period as a function of trial for each group and examined the function which best described each (Table S2). These findings also echoed our behavioural characterisation. In replete rats, dopamine responses were generally weak and varied minimally across trials, with R100 responses best described by linear regression and R450 responses best described by a sigmoid capturing a weak dopamine response on early trials which rapidly faded. D100 dopamine responses were best fit by linear regression, reflecting a gradual decrease in evoked dopamine magnitude which nevertheless remained positive throughout the session. For D450 rats, evoked dopamine release was best described by a sigmoid, reflecting robust dopamine release that persisted beyond early trials before declining later in the session (trial of transition midpoint (x_0_) from sigmoid fit = 17). The transition midpoint for D450 rats occurred after infusion of ∼3.1 milliequivalents of sodium, supporting the notion that the step-like change in dopamine is due to the development of post-ingestive satiating signals (Krause et al., 2010; Roitman et al., 1997) Together, these results demonstrate that sodium-evoked dopamine release is dynamic within session and is impacted by ongoing ingestion.

Finally, we examined whether sodium-evoked dopamine release could reliably distinguish rats with varying physiological needs. To do so, we first computed the euclidean distance between pairwise comparisons of rats based on their dopamine signal across all infusion periods (see Methods; Figure 2G and S3C). Next, we used multidimensional scaling to visualise this dissimilarity structure in two dimensions (Figure 2H). This analysis revealed that rats within each condition were generally more similar (i.e., closer) to one another than rats in other conditions. R100 and R450 rats were similar to one another, while D100 rats differed greatly from all other groups. The cluster of D450 rats fell in between these two extremes, consistent with a transition in need state from deplete to replete across trials reflected in the dopamine responses of these rats. Quantifying these observations, we found that a cross-validated classifier could identify the group to which rats belonged based on their sodium-evoked dopamine release across trials with accuracy well above chance (Accuracy = 45.0%; Chance = 25.0%; binomial test *p* = 0.005). Classifier confusions were most common between the replete groups, reflecting the poorer separation between R100 rats versus R450 rats, while deplete groups were classified more reliably (Figure 2I). Together, these results demonstrate that sodium-evoked phasic dopamine release reliably reflects physiological need, even as that need wanes with ingestion.

### Changes in sodium-evoked dopamine release precede the shift in behavioural expression during satiation

So far, we have shown that sodium-evoked dopamine release and appetitive behavioural expression reflect physiological need in commensurate ways. In D450 rats, both dopamine release and behaviour exhibited a step-like transition across the session, from high dopamine release and appetitive behaviour early in the session to low dopamine release accompanied by reduced appetitive behaviour later in the session. Interestingly, while both dopaminergic and behavioural assays were best fit by a sigmoid in these rats (Figure 1F and 2F), the point of transition differed between the two with the average dopamine transition point (trial 17) preceding the average behavioural transition point (trial 24). This discrepancy suggests that changes in sodium-evoked dopamine release reflecting satiation might precede changes in sodium-evoked appetitive behaviour. If so, then we should observe a consistent offset between the change in sodium-evoked dopamine release and appetitive behaviour in individual rats.

To test this possibility, we crosscorrelated the trial-wise dopamine signal (AUC of infusion period) with the trial-wise probability of appetitive behaviour in individual rats, varying the lag between the two (Figure 3A-C). Consistent with our group-level finding, we found that the lag which produced the highest correlation between the dopamine and behavioural signals reliably differed from zero across rats (one-sample t-test vs. zero, *t*_8_ = 2.79, *p* = 0.023), with dopamine preceding behaviour (median optimal lag: 3 trials). When repeating this analysis for rats in other groups the optimal lag did not significantly differ from zero (all *p*s > 0.05), demonstrating that this finding is specific to the dynamics associated with satiation present in D450 rats. Moreover, the optimal lag produced numerically high and statistically higher correlations in the D450 rats when compared with rats in other groups (Figure 3C; one-way ANOVA; *F*_3,35_ = 10.29, *p* < 0.001), reinforcing the unique alignment between neural and behavioural signals in this cohort. Together these findings provide corroborative evidence that physiological need-dependent dopamine dynamics precede behavioural changes associated with revaluation during sodium satiation.

**Figure 3.**
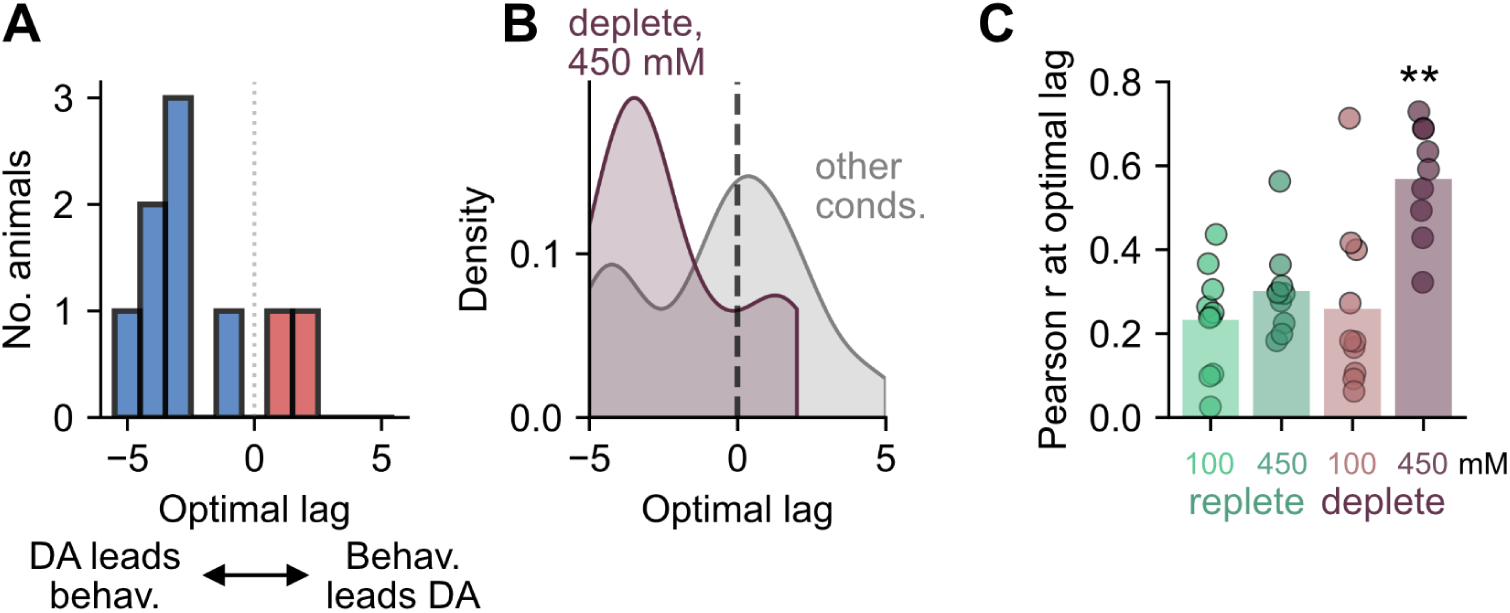
Behavioural and dopamine responses to NaCI show high correlation in rats experiencing a hedonic switch. (A) Optimal lags after crosscorrelating appetitive behaviour and dopamine responses across the session for rats receiving 450 mM NaCI in the deplete state (D450 rats). Blue bars show negative lags, red bars show positive lags. (B) Kernel density estimation plot showing optimal lags from rats in (A), relative to rats in all other conditions. Plot showing Pearson r values at the optimal lag for each condition. Bars are mean and circles are individual rats. **, *p* < 0.01 vs. all other groups.

### Accounting for individual differences reveals sharp transition in dopamine release and behaviour

As noted above, D450 rats exhibited on average a transition from sodium-evoked high dopamine/high appetitive behaviour to a low dopamine/low appetitive behaviour as trials proceeded (Figure 4A and 4E). While this transition was relatively gradual for both measures (k_behaviour_ = 0.31; k_dopamine_ = 0.19), inspection of the data from individual rats suggested that transitions occurred abruptly within individuals but varied in the trial at which the transition occurred (Figure S3). Given this observation, we next tested whether dopamine release and behavioural expression changed more abruptly across trials in individual D450 rats than expected based on the group mean data.

**Figure 4.**
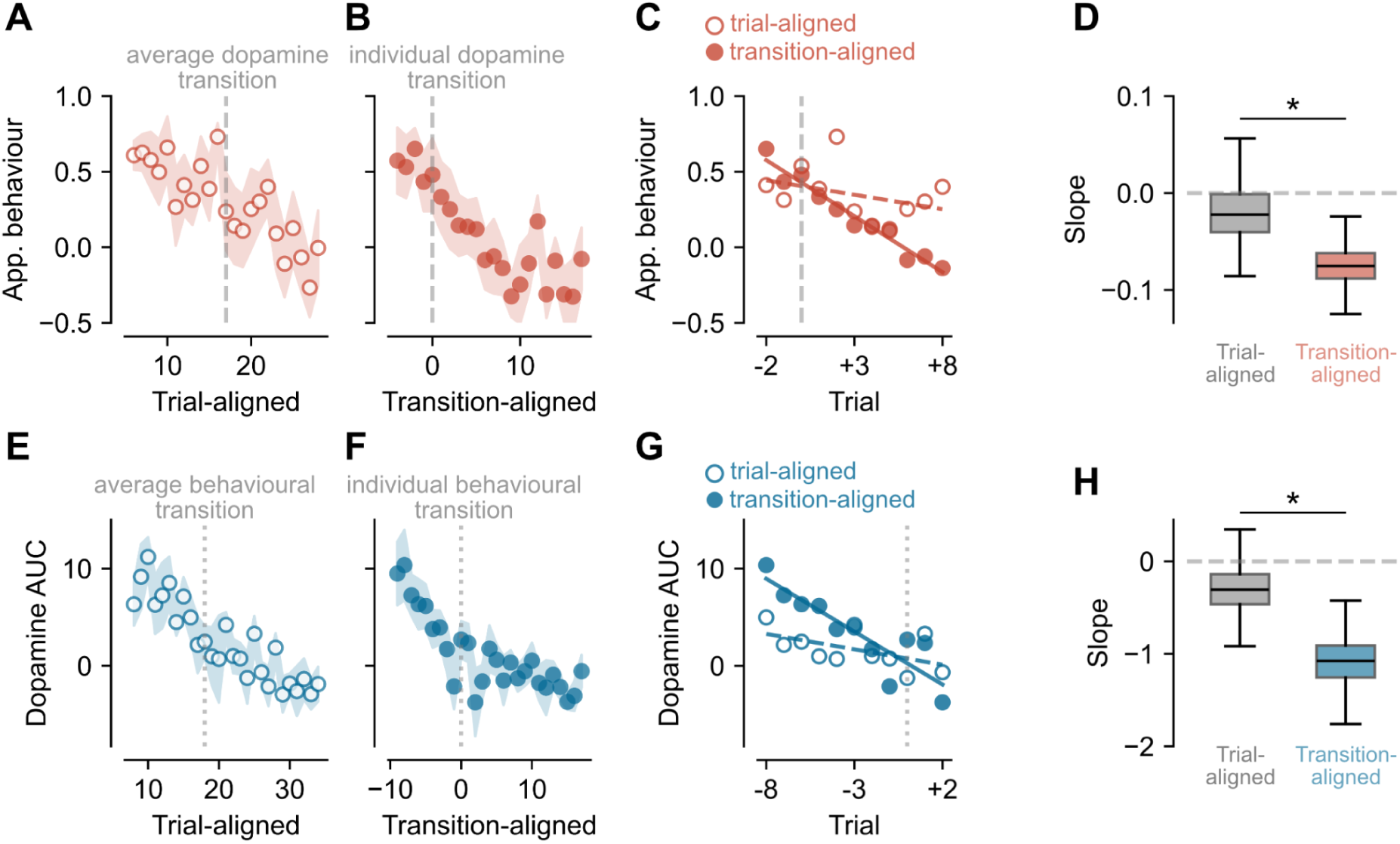
Realignment of responses per individual rat sharpens the hedonic and dopamine transitions. (A) Plot showing behavioural responses to NaCI infusions across trials in deplete rats receiving 450 mM NaCI. Circles are mean and shaded area is SEM. Grey dashed line is mean dopamine transition. Data are trimmed to include the same number of trials in A and B. (B) Similar plot to (A) but after realignment to each rat’s individual dopamine transition point as calculated from velocity of changing dopamine response. Grey dashed line is individual dopamine transition for each rat. Only trials with data for all rats are included. (C) Behavioural responses for 5 trials before and 5 trials after transition plus lag from crosscorrelation (+3 trials). Linear regression shown for trial-aligned data (open circles, dashed line) and transition-aligned data (closed circles, solid line). (D) Boxplot showing slope values for trial-aligned and transition-aligned data after 1000 bootstrapped comparisons. *, *p* < 0.05. (E-H) As in A-D but for dopamine responses aligned to behavioural transitions. Dotted grey line in E-G indicates behavioural transition point (average or individual). G shows 5 trials before and 5 after transition minus lag from crosscorrelation (−3 trials).

To test this possibility, we compared the change in the original behavioural and dopamine signals (i.e., trial-aligned) to the change observed when each D450 rat’s data were realigned according to its individual-specific transition trial (i.e., transition-aligned; Figure 4B and 4F). For each rat, we defined the transition trial as the trial during which the dopamine or behavioural signal changed most rapidly. To ensure that this transition trial selection did not bias our outcomes, when realigning the behavioural signal we chose the dopamine transition trial adjusted by the median optimal lag (+3 trials; from Figure 3A), and when realigning the dopamine signal we chose the behavioural transition trial adjusted by the median optimal lag (−3 trials). Next, to estimate the rate of change, we fit a linear slope to the ±5 trials flanking the lag-adjusted transition trial, and compared the naive trial-aligned slope to the transition-aligned slope (Figure 4C, 4D, 4G, 4H). Using a bootstrapping procedure to estimate the effect of aligning to individual transitions, analysis showed that for both the appetitive behavioural signal and the dopamine signal, transition alignment yielded significantly steeper slopes, relative to the original trial-aligned data (Figure 4D and 4H; paired bootstrapped-difference test with *n* = 1000 bootstrap resamples; behaviour: *p* = 0.039; dopamine: *p* = 0.014). Transition-aligned slopes also significantly differed from zero (behaviour: *p* < 0.001; dopamine: *p* < 0.001). Together, these results demonstrate that satiation-like changes in these signals occurred more abruptly in individual rats than would be expected from the group-level changes alone.

### Infusion-evoked dopamine release and behavioural expression track cumulative sodium consumption, not infusion number or volume

We have demonstrated that physiological need modulates sodium-evoked dopamine release and behavioural expression. These effects are seen both when need state is manipulated prior to infusions via sodium depletion, and during ongoing ingestion especially in D450 rats. Interestingly, D100 rats received the same number of trials and volume of fluid yet exhibited a much more modest change in dopamine and behaviour, relative to D450 rats. This could suggest that dopaminergic and behavioural changes track the cumulative amount of sodium consumed (i.e., the number of sodium ions) rather than the number or volume of infusions.

To test this possibility, we compared infusion-evoked dopamine release and behavioural expression between D100 and D450 rats in two ways: once when the data were organised naively as a function of trial number and again when the data were organised as a function of the cumulative amount of sodium delivered during each session. As described previously, when data were organised by trial number, we observed a significant difference between D100 and D450 rats in both behavioural expression and dopamine release (Figure 5A and 5D; two-way mixed ANOVA main effect of Concentration; behaviour: *F*_1,18_ = 34.47, *p* < 0.001; dopamine: *F*_1,18_ = 36.56, *p* < 0.001). However, when the data for the two groups were organised by cumulative sodium, we observed striking similarity between the groups in both behavioural expression and dopamine release (Figure 5B, 5C, 5E and 5F). Reflecting this similarity, no significant effects of Concentration (OLS with cluster-robust standard errors; *β* = −0.24, *p* = 0.208), sodium delivered (*β* = −0.10, *p* = 0.629), or an interaction (*β* = −0.16, *p* = 0.481) were observed when comparing behavioural expression. When comparing dopamine, a significant main effect of Concentration (*β* = 7.13, *p* = 0.040), sodium delivered (*β* = −17.12, *p* < 0.001), and a significant interaction (*β* = 12.77, *p* = 0.013) were observed. Inspection of the replotted data suggested that these differences were driven by large dopamine responses to sodium infusions early in the session in D100 rats. To assess this, we re-ran the analysis after removing the first ∼30 mg sodium delivered (first 13 trials for D100 rats and first 3 trials for D450 rats). After this adjustment, there was only an effect of the amount of sodium delivered (*β* = −12.38, *p* = 0.010), but no effect of Concentration (*β* = 6.09, *p* = 0.168) or any interaction (*β* = 8.15, *p* = 0.094). These similarities indicate that our neural and behavioural measures index satiation as a function of the amount of sodium delivered and not other correlated measures such as number or volume of infusions. Moreover, these results provide an explanation for the lack of satiation-dependent changes in D100 rats, namely that these rats do not consume a sufficient amount of sodium (e.g., only ∼2 mEq infused relative to ∼2.5-3.0 mEq lost) to be satiated in the course of an experimental session.

**Figure 5.**
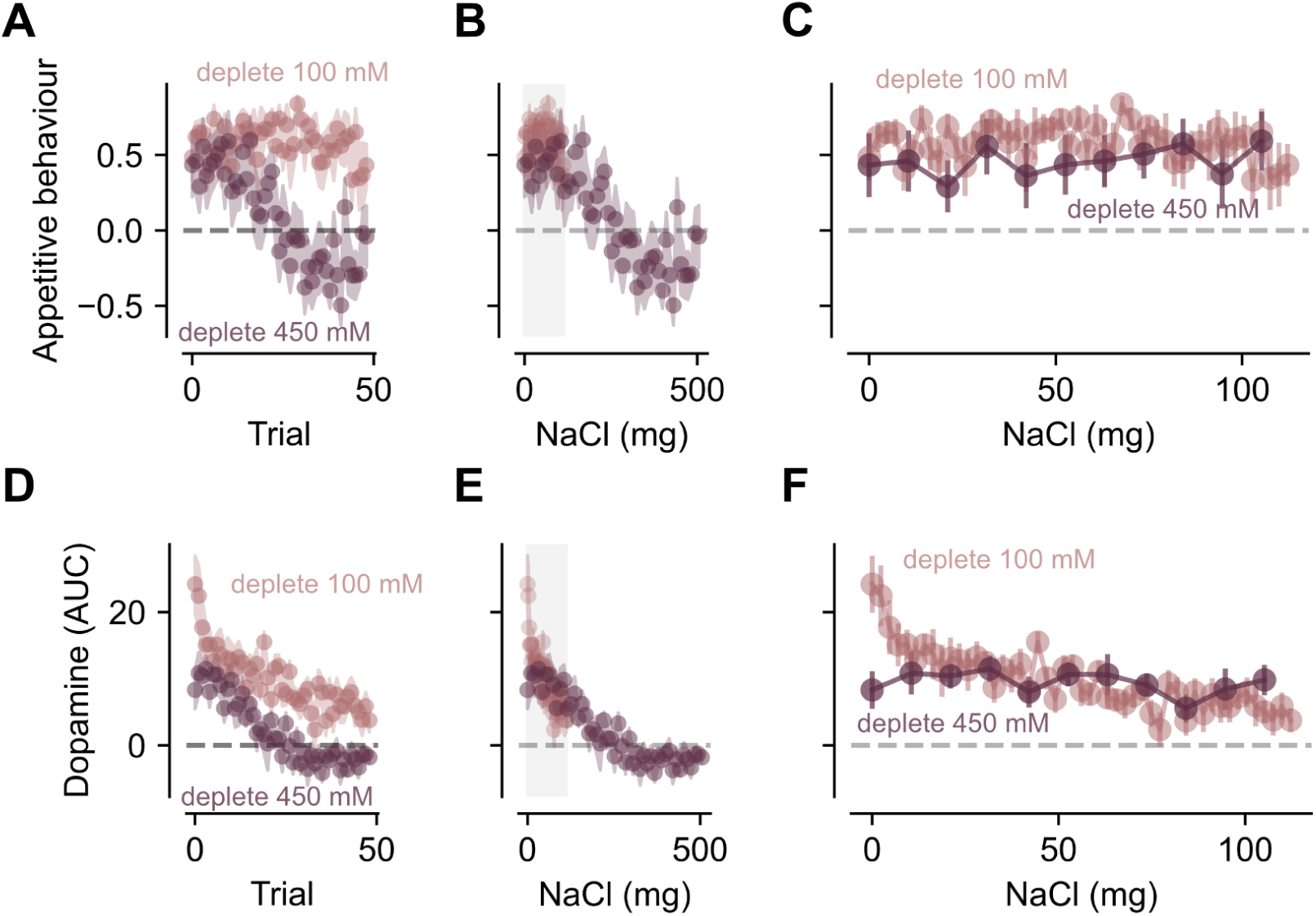
Amount of salt ingested is integrated by behaviour and dopamine to determine need state. (A) Plots showing appetitive behaviour as a function of trial for deplete 100 mM and deplete 450 mM rats. Circles are mean and shaded area is SEM. (B) Plots showing appetitive behaviour as a function of NaCI delivered across the session for deplete 100 mM and deplete 450 mM rats. Circles are mean and coloured shaded area is SEM. Grey shaded area shows area expanded in C. (C)Expanded plot showing rats receiving 100 mM and 450 mM NaCI as a function of NaCI delivered. Circles are mean and error bars show SEM. (D-F) As in A-C but for dopamine AUC responses.

## Discussion

Sodium appetite is an innate, selective behavioural response to sodium deficit (Krause & Sakai, 2007; Richter, 1936) and provides a classic model for understanding how physiological need shapes motivated behaviour. Indeed, sodium depletion switches the behavioural response to concentrated sodium solutions from avoidance/rejection to approach/avid consumption (Berridge et al., 1984; Robinson & Berridge, 2013). The mesolimbic dopamine system promotes and reinforces goal-directed actions (Stuber, 2023). In particular, phasic dopamine release, which arises from bursting activity of dopamine neurons, is thought to be crucial for these functions. Consistent with these frameworks, and similar to other drivers of ingestive behaviour (e.g., caloric/water restriction; Hsu et al., 2020; Reichenbach et al., 2024), mesolimbic dopamine signalling is recruited by the taste of sodium but only after sodium depletion (Cone et al., 2016; Fortin & Roitman, 2018; Ozawa et al., 2025). Here, we show that ongoing ingestion of sodium ‘derecruits’ mesolimbic dopamine signaling. Moreover, steep reductions in taste-evoked dopamine precede the transition away from appetitive behaviour in the sodium deplete rat. Our results give critical insight into how satiation signals influence neural circuits underpinning motivated behaviour.

### Behavioural responses to sodium

Sodium depletion radically changes behaviour directed at sodium-containing solutions. Prior work has shown that sodium depleted rats select solutions with cation specificity and regardless of the anion (Breslin et al., 1993). Sodium deplete rats will readily lever press to self-administer sodium (Quartermain et al., 1967) and develop Pavlovian approach responses for sodium-predictive cues, even after only experiencing pairings of cues with sodium while replete (Robinson & Berridge, 2013). In replete rats, behavioural responses to brief intraoral infusions of high concentration sodium elicit a behavioural pattern consistent with aversion and these responses switch to appetitive following depletion (Berridge et al., 1984). Our behavioural findings are consistent with this literature. Here, deplete rats exhibited appetitive behaviour to both low and high concentrations of sodium whereas such behaviour was wholly absent in replete rats.

When given free access to sodium, deplete rats rapidly restore lost sodium within minutes, after which approach and consumption ceases (Roitman et a. 1997). Treatment with the diuretic furosemide causes approximately 2.5 mEq loss of sodium in the urine and rats typically overconsume sodium relative to loss (Krause et al., 2010; Lundy et al., 2003; Rowland & Fregly, 1992). Using intraoral delivery, we were able to control the rate of repletion by adjusting the number of trials and the concentration of sodium. As such, delivery of 100 mM NaCl over fifty trials (2.0 mEq) was insufficient to replenish lost sodium and appetitive behaviour remained high throughout the session. In contrast, 450 mM NaCl infusions delivered more sodium (9.0 mEq) than lost. The striking and novel result of this approach is that, using a deep learning-based pose estimator (DeepLabCut) together with a behavioural classifier (SimBA), we quantified the trial-by-trial change in appetitive behaviour and were able to identify satiation transition points for individual rats.

### Dopamine responses to sodium

The biological architecture for recruitment of mesolimbic dopamine by the taste of sodium following depletion has been well established. Sodium depletion initiates a cascade of hormone release that includes elevations in circulating angiotensin II and aldosterone. These hormones act on central circuits to organise sodium appetite (Sakai et al., 1986). As such, angiotensin II binds to receptors within the lamina terminalis, a set of structures that includes the subfornical organ (SFO). Activation of subsets of SFO neurons is sufficient to drive water consumption (Nation et al., 2016; Oka et al., 2015) and recruit phasic dopamine responses to water (Grove et al., 2022), sodium (Zhang et al., 2023), and cues predicting fluid (Hsu et al., 2020). While the SFO does not project directly to mesolimbic circuitry, it may recruit dopamine neurons via connections with the lateral hypothalamus (Hurley et al., 2018). Aldosterone, on the other hand, acts on nucleus of the solitary tract (NTS) neurons in the hindbrain that express 11β-hydroxysteroid-dehydrogenase (HSD2) to drive sodium appetite (Gasparini et al., 2024; Jarvie & Palmiter, 2017; Resch et al., 2017). These NTS neurons form an ascending circuit that may directly (Geerling & Loewy, 2008) or indirectly (Hurley et al., 2018; Shin et al., 2011) recruit mesolimbic dopamine neurons. Moreover, we have shown that phasic dopamine responses to sodium (and sodium-predictive cues) are only engaged when animals are deplete and require the ability to detect the taste of sodium in solution (Cone et al., 2016; Fortin & Roitman, 2018). Thus, the mechanisms underlying sodium-evoked recruitment of mesolimbic dopamine is well described. In contrast, in the current work, we focus on the complementary process of “derecruitment” of dopamine as sodium balance is restored.

Our principal finding is that phasic dopamine responses to sodium decrease as deplete rats restore their sodium levels. In this light, a key question is how sodium ingestion is relayed to dopamine neurons. The aldosterone-sensitive HSD2 neurons in the NTS project, in part, to the pre-locus coeruleus (pre-LC; Gasparini et al., 2019) and pre-LC activity is elevated after sodium depletion. Upon encountering sodium, the activity of pre-LC neurons drops precipitously within seconds (Lee et al., 2019). Similar to our prior work measuring dopamine responses (Cone et al., 2016; Fortin & Roitman, 2018), changes in pre-LC activity are dependent on the presence and detectability of the sodium ion. Decreasing activity in pre-LC could be the signal for reduced sodium drive and account for the decreasing dopamine responses observed in our current experiment. Yet such an account is insufficient to explain our data in full. Pre-LC activity decreases regardless of sodium concentration and in response to just seconds of access (Lee et al., 2019). In our data, the rate of dopamine decrease was best described by the absolute amount of sodium consumed. Moreover, transitions in dopamine and behaviour occurred only after the amount of sodium consumed exceeded what is typically lost suggesting a post-oral component of negative feedback. Indeed, studies using sham drinking show that oral stimulation of sodium taste alone is insufficient to satiate sodium appetite (Krause et al., 2010; Roitman et al., 1997). As mentioned above, SFO is another crucial node that modulates sodium consumption and changing activity here could be responsible for decreases in dopamine responses during satiation. In fact, recent work identified a subset of SFO neurons that mediate tolerance for the taste of high concentration sodium in the deplete state (Zhang et al., 2023). Most interestingly, activation of these neurons recruited a dopamine response to the taste of high concentration sodium. It is unknown how their activity is modulated during ingestion but this work intriguingly suggests that the reduction in dopamine and appetitive responses we observed are underpinned by a reduced tolerance for high concentration sodium.

Phasic dopamine responses decreased only after substantial sodium was infused suggesting post-oral, post-ingestive negative feedback. The “overshoot” in sodium ingested versus sodium lost observed by us and others also hints that a slower hormonal signal may be necessary to convey repletion. The gut hormone secretin was recently linked to the generation of sodium appetite (Liu et al., 2023). Indeed, secretin receptors in the NTS appear necessary for depletion-induced sodium appetite and the activity of NTS secretin receptor-expressing neurons is elevated in the minutes after intravenous secretin. As sodium channels are expressed throughout the gut and in the kidneys (Akopian et al., 1997; Lara et al., 2012), accumulating sodium could lead to a decrease in secretin, angiotensin II, and aldosterone production and represent a more slowly changing signal to eliminate central drive.

### Rapid switching of behaviour and dopamine

There was no difference in initial appetitive behaviour between D100 and D450 rats. This may reflect a ceiling effect in our measure as the dominant component of our appetitive behaviour index was a decrease in movement. While our measure of appetitive behaviour did not capture initial differences between D100 and D450 rats, the magnitude of dopamine responses was initially higher in D100 rats. Rats prefer near-isotonic (∼150 mM) concentrations of sodium relative to hypertonic even after sodium depletion (Breslin et al., 1993). Our dopamine results are therefore consistent with literature supporting a role for phasic dopamine in carrying information related to subjective preference (Schultz, 2015). Indeed, through the first 13 trials, dopamine release was higher in magnitude for D100 rats even after accounting for the amount of sodium infused.

Dopamine and appetitive behaviour transitioned from high to low states in a step-like manner for D450 rats as infusions progressed. The decrease in dopamine and appetitive behavior became steeper as we accounted for individual differences in transition points. Importantly, decreases in dopamine reliably preceded decreases in appetitive behaviour by approximately 3 trials. These data are consistent with reinforcement learning (RL) models where phasic dopamine is critical for updating the value of stimuli (Fraser et al., 2025) and actions (Gershman & Uchida, 2019). Importantly, most RL models fail to account for changing internal state. Recent iterations (Duriez et al., 2023; Yoshida et al., 2025) though, highlight the importance of drive states. More recently, arguments have been made that phasic dopamine is crucial for signaling reward magnitude which, in turn, influences the learning rate in RL models (Gong et al., 2026). While increasing drive has been repeatedly shown to increase the magnitude of dopamine responses to needed stimuli (Roitman & McCutcheon, 2025), the present data highlight that decreases in drive may be just as important for attenuating phasic dopamine responses. In this framework, steeply diminishing dopamine events from trial to trial likely lead to rapid devaluation of ongoing ingestion and thus a decrease in appetitive behaviour.

## Resource availability

All data are available on Zenodo at https://doi.org/10.5281/zenodo.21394487 and analysis code is available at https://github.com/jaimemcc/bazzino.

## Acknowledgements

We gratefully acknowledge the mentorship and guidance of Dr. Ted Hsu and Dr. Jamie Roitman throughout this project. We also thank Dr. Ted Hsu, Dr. Max Loh, Dr. Vaibhav Konanur, and Dr. Rachel Donka for their invaluable technical support and training in fibre photometry data collection and analysis. This work was supported by funding from National Institutes of Health (DA025634 to MFR) and the Tromsø Research Foundation (19-SG-JMcC to JEM).

## Author contributions

Conceptualization: PB, MFR; Formal Analysis: PB, DCR, ATK, JEM; Funding Acquisition: MFR, JEM; Investigation: PB; Methodology: PB, ATK, MFR; Supervision: MFR, JEM; Writing - Original Draft: PB, ATK, MFR, JEM; Writing - Review & Editing: DCR.

## Declaration of generative AI and AI-assisted technologies in the writing process

No generative AI has been used in the writing process.

## STAR★Methods

### Key resources table

| Reagent/Resource | Source | Identifier |
| --- | --- | --- |
| **Antibodies** |  |  |
| Chicken polyclonal anti-GFP | Abcam | Cat#AB13970;<br>RRID:AB_300798 |
| Donkey polyclonal anti-chicken AF488 conjugated | Jackson Immunoresearch | Cat#703-545-155;<br>RRID:AB_2340375 |
| **Bacterial and virus strains** |  |  |
| pAAV-hSyn-GRAB_DA1h | Addgene / Sun et al., 2018 | Cat#113050-AAV9;<br>RRID:Addgene_113050 |
| **Chemicals, peptides, and recombinant proteins** |  |  |
| Furosemide | Sigma-Aldrich | Cat#F4381 |
| **Deposited data** |  |  |
| Behavioural and photometry data | Zenodo | https://doi.org/10.5281/zenodo.21394487 |
| Experimental models: Organisms/strains |  |  |
| Rat: Long-Evans (LE) | University of Illinois Chicago animal unit | RRID:RGD_60991 |
| Software and algorithms |  |  |
| JASP (v0.97) | JASP | RRID:SCR_015823 |
| Other |  |  |
| Sodium deficient diet | Envigo | Cat#TD 90228 |
| Fibre optic cannulas (flat 400 μm core, 0.48 NA) | Doric Lenses | Cat#: MFC_400/430-0.48_10.0mm_ZF2.5_FLT |
| RZ5P | Tucker Davis Technologies | RRID:SCR_024878 |
| Visible femtowatt photoreceiver | Newport | Cat#2151 |

### Animals

Adult male (n = 10) and female (n = 10) wild-type Long Evans rats (> 270 g) were maintained on a 12-hour light-dark cycle (lights on at 7:00am) in a temperature- and humidity-controlled environment. Rats had ad libitum access to water and standard laboratory chow unless otherwise stated. All experiments were conducted during the light cycle. Animal care and use was in accordance with the National Institutes for Health Guide for the Care and Use of Laboratory Animals and approved by the Institutional Animal Care and Use Committee at the University of Illinois at Chicago.

### Surgery

Rats were anesthetised via isoflurane inhalation [SomnoSuite; 4% (vol/vol) at 500 mL/min for induction and thereafter maintained at 2% (vol/vol) with the infusion rate dependent on the rat’s weight] and the head secured in stereotaxic frame (Kopf). Subcutaneous analgesia (0.1 mL of 5 mg/mL Meloxicam) was provided before making any incision to the animal as well as for two days following surgery. For the recording of dopamine release in the nucleus accumbens (NAc) lateral shell (AP: 1.5 mm, ML: 2.5 mm, DV: −8.0 mm virus/-7.9 mm fibre optic, relative to bregma), 1 uL of a genetically encoded GPCR-activation-based-dopamine fluorescent sensor (pAAV-hSyn-GRAB_DA1h, Addgene; Sun et al., 2018) was unilaterally injected at a rate of 0.1 µL/min. Then, an optic fibre (flat 400-µm core, 0.48 numerical aperture [NA], Doric Lenses Inc.) was immediately implanted 0.1 mm above the injection site. An intraoral catheter was also implanted. In brief, a length of polyethylene (PE) tubing (Scientific Commodities, Inc) was flanged at one end and a Teflon washer placed over it. The other end was mated to a 19-gauge stainless steel needle (Component Supply). The needle was inserted just lateral to the first maxillary molar, exteriorized through the head incision and fixed, along with the fibre optic, to skull screws with dental acrylic. Catheters were flushed daily with deionized water for the remainder of the experiment to habituate rats to intraoral delivery and to prevent clogging. Rats were given two weeks to recover from surgery and for virus expression.

### In vivo fibre photometry

In vivo fibre photometry was used in awake and behaving rats to record dopamine release in the NAc lateral shell (Figure 1A). The light-emitting diodes (LED) used to excite GRAB-DA1h were 465 nm (dopamine-dependent) and 405 nm (dopamine-independent). Intensity of the 465 nm and the 405 nm light was sinusoidally modulated at 211 and 532 Hz, respectively. The emitted GRAB-DA fluorescence was then recorded via the chronically implanted optical fibre into an optical fibre patch cord and transmitted to a photoreceiver (Newport, Model 2151). Data acquisition and demodulation was managed by Tucker Davis Technologies (TDT, Model RZ5P; Synapse Suite). Key behaviourally relevant events were transmitted as time-stamped TTLs to the RZ5P and recorded in software (Synapse Suite, TDT).

### Behavioural procedures

Two weeks following surgery, rats were habituated to standard operant chambers (in cm, L: 29.5, W: 31.8, H: 27.4; ENV-009A-CT, Med Associates Inc.) for two days. During each habituation day, rats were administered one session of 30 trials, with each trial consisting of a 5 s intraoral water infusion (200 µL) and a 35-55 s inter-trial interval (interval duration selected randomly on each trial). Infusion onset and offset were controlled by a solenoid valve. A reservoir containing fluid was positioned above the solenoid and gravity-fed into the solenoid. A length of tubing from the other solenoid port was connected to the intraoral catheter. On test days, rats received 10 s intraoral infusions (400 µL) of either a hypotonic (100 mM) or hypertonic (450 mM) NaCl solution for a total of 50 trials (intertrial intervals as in habituation sessions). Dopamine recordings were made under sodium replete and deplete states.

### Sodium depletion

Sodium appetite was induced in the laboratory by injecting two, equal-volume subcutaneous injections, spaced one hour apart, of the diuretic furosemide (20 mg/mL, 0.5 mL/kg; Sigma Aldrich). Diuresis was confirmed by weighing the animals before and two-hours following the first injection. Rats were considered sodium deplete if they lost at least 15 g of body weight in the two-hour period. Before injections, food and water were removed. Following the 2-hours after the first injection, rats were maintained for 24 h on distilled water and a sodium deficient diet (TD 90228, Envigo).

### Immunohistochemistry

Following experiment completion, rats were anesthetized with sodium pentobarbital (100 mg/kg) and transcardially perfused with 0.01 M phosphate-buffered saline (PBS) followed by 10% buffered formalin solution (HT501320, Sigma Aldrich). The brains were removed and stored in 20% sucrose in formalin for 24 h and then sectioned at 30 µm on a freezing stage microtome (SM2010R, Leica Biosystems). Sections were processed to label for green-fluorescent protein (GFP) as an indicator of GRAB-DA1h. Tissue was incubated in primary (chicken anti-GFP; AB13907, Abcam) and secondary (AF488 conjugated donkey anti-chicken; Jackson Immunoresearch) antibody overnight at 4°C. Sections were then cover slipped on slides for visualization. All images were collected using a BX43 Olympus microscope with Lumen Dynamics for visualising fluorescence and a XM10 monochrome camera. Only data from rats with confirmed GFP expression and fibre optic placement in the NAc were analysed.

### Pose estimation and behavioural classification

Behavioural responses to NaCl infusions were measured during different sodium need states and analyzed using artificial intelligence-based DeepLabCut (DLC), an open-source markerless pose estimator based on deep neural networks (Mathis et al., 2018). A video camera (Wo-We, 10 frames/second with 480 x 640 pixels resolution) was positioned at the top of the behavioural chamber.

We randomly split 95% of the data for training the model and the rest for testing. We trained the model on 230 frames from 31 videos, (∼8 frames/video sampled in DLC using the k-means clustering approach annotated by one human). We annotated twelve body parts of interest (nose, left ear, right ear, head base, back 1-4, tail base, tail 1-3, and tail tip). We trained the model with 200,000 iterations and achieved train and test RMSE 1.56 and 7.42 pixels, respectively. With a detection confidence cutoff (*P*_cutoff_ = 0.6), the test RMSE decreased to 6.28 pixels. Train mAP/mAR were 100%; test mAP/mAR were 93.17% and 93.33%.

To identify appetitive behaviour, we used SimBA (Simple Behavioural Analysis) (Goodwin et al., 2024). For feature extraction we used a subset of body parts tracked by DLC (left ear, right ear, nose, head base, tail base), filtered in SimBA using interpolation (“body-parts: nearest” option) and gaussian smoothing, and wrote a custom feature extraction script (https://github.com/jaimemcc/bazzino/blob/main/src/hybrid_feature_extractor.py). We used the low concentration (100 mM NaCl) deplete rats (D100) as a positive control and assigned all video frames when these rats received intraoral infusions as containing “appetitive” behaviour. We used these assignments to train a model to identify appetitive behaviour in rats in other groups. SimBA uses a random forest classifier and the hyperparameters we used were: criterion = Gini impurity, max features = square root, minimum sample leaf = 1, and number of estimators = 2,000. To identify appetitive behaviour in the low concentration deplete group, we used a “leave-one-out” validation where separate models were trained that did not include data from the rat to be tested.

### Data analysis and statistical methods

We used custom Python scripts to perform all analyses (Python, version 3.12, Python Software Foundation, Wilmington, DE) with the exception of mixed effects ANOVA, which were performed with JASP (v0.97). Data wrangling, plotting, and statistical analysis used the following packages and libraries: Numpy (Harris et al., 2020), Pandas (McKinney, 2010), Matplotlib (Hunter, 2007), Seaborn (Waskom, 2021), and Scipy (Virtanen et al., 2020). Analysis code is deposited at https://github.com/jaimemcc/bazzino.

For behavioural data, to obtain a value representing the probability of appetitive behaviour on any given trial, we constructed a null distribution based on repeated shifting of the appetitive probability vector by a fixed amount and excluding overlap with real infusions. We used this null distribution to derive the median probability of appetitive behaviour across the whole session and then for each infusion we calculated the difference between bins (frames) > median and bins < median. Alternatively, this can be expressed as *2p-1*, where p is the proportion of bins during the infusion that exceed the median. This probability ranges from −1 (all infusion bins < median) to +1 (all infusion bins > median) and was used for subsequent trial-by-trial analyses.

For fibre photometry data, the signal was corrected for movement artifacts and bleaching using a Fourier-based subtraction of the 405 nm (dopamine-independent) signal from the 465 nm (dopamine-dependent) signal in the frequency domain (Konanur et al., 2020). The corrected signal was aligned to the first 49 solenoid openings, downsampled to 10 Hz, and converted into a matrix of size 49 x 200, representing 5 s before and 15 s after solenoid opening. Each trial was z-scored using the mean and standard deviation of the 5 s baseline. For trial-by-trial comparisons, the AUC of the 10 s infusion period was used.

For analysis of mean appetitive behaviour and mean dopamine AUC across the whole session, each of the four subgroups were compared against zero using two-sided Wilcoxon signed-rank tests (non-parametric one-sample paired-rank). In addition, a linear mixed-effects omnibus model was used with Condition (deplete vs. replete), Concentration (100 mM versus 450 mM), and Sex (male vs. female) as factors. No main effect of Sex or significant interactions were found so sexes were pooled for further analyses. Significant interactions were followed up with Holm-adjusted post hoc tests.

To determine whether the group identity of rats could be determined by either their behavioural or dopamine responses across trials, we found the euclidean distance between individuals in multidimensional space and plotted the average distance between individuals within each group. We used multidimensional scaling to project distances between individual rats and group averages onto a two-dimensional space. To test whether group identity could be accurately predicted from either behavioural or dopamine responses, we used a logistic regression-based classifier via leave-one-out method and tested the predictions versus chance using a binomial test.

In high concentration deplete rats (D450), we identified the trial number at which behaviour or dopamine transitioned from high to low using the maximum negative velocity of the derivative after smoothing data (gaussian, sigma=1). One rat’s behaviour did not change across the session and could not be fit with a sigmoid, so was excluded from analysis requiring realignment to transition points.

To analyse behaviour and dopamine for deplete rats (D100 and D450) as a function of sodium delivered, we fit ordinary least squares regression models with amount of sodium (z-scored, continuous), Concentration (100 mM versus 450 mM), and their interaction as predictors.

Subject-clustered robust standard errors (clustered by mouse), were used to handle repeated observations within subjects.

