## Supplemental Material for "Rapidly decreasing phasic dopamine responses precede a hedonic switch during satiation of sodium appetite"

Supplemental material for Bazzino et al.

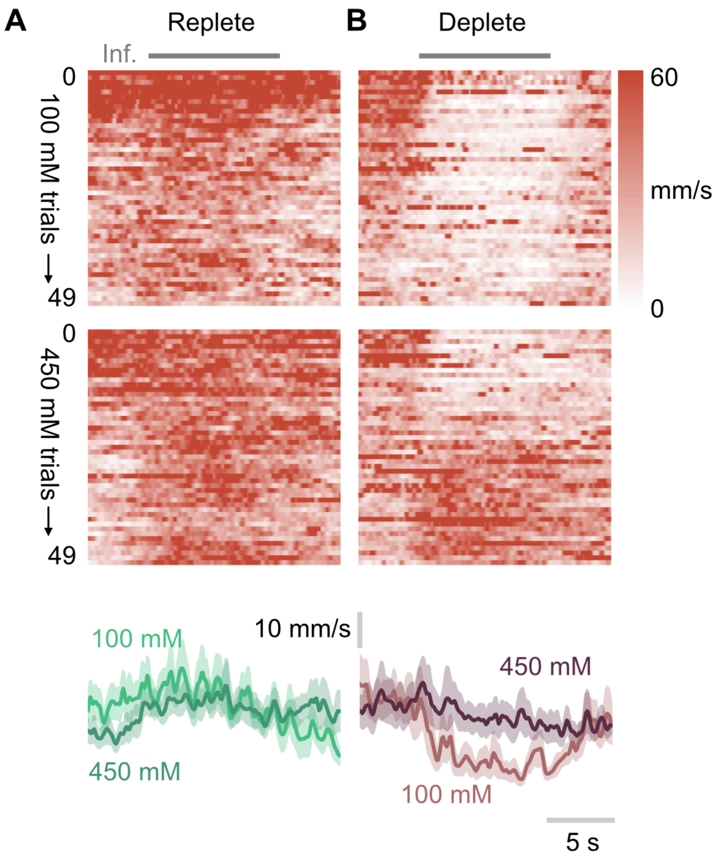


**Figure S1. Head movement during intraoral infusions of NaCl under different physiological conditions.**

(A) Heatmaps showing movement of the head after intraoral infusions of low (100 mM; upper heatmap) and high (450 mM; lower heatmap) NaCl. Bottom panels show mean ± SEM. n=10 rats in each group.

(B) As in (A), but for rats after sodium depletion.

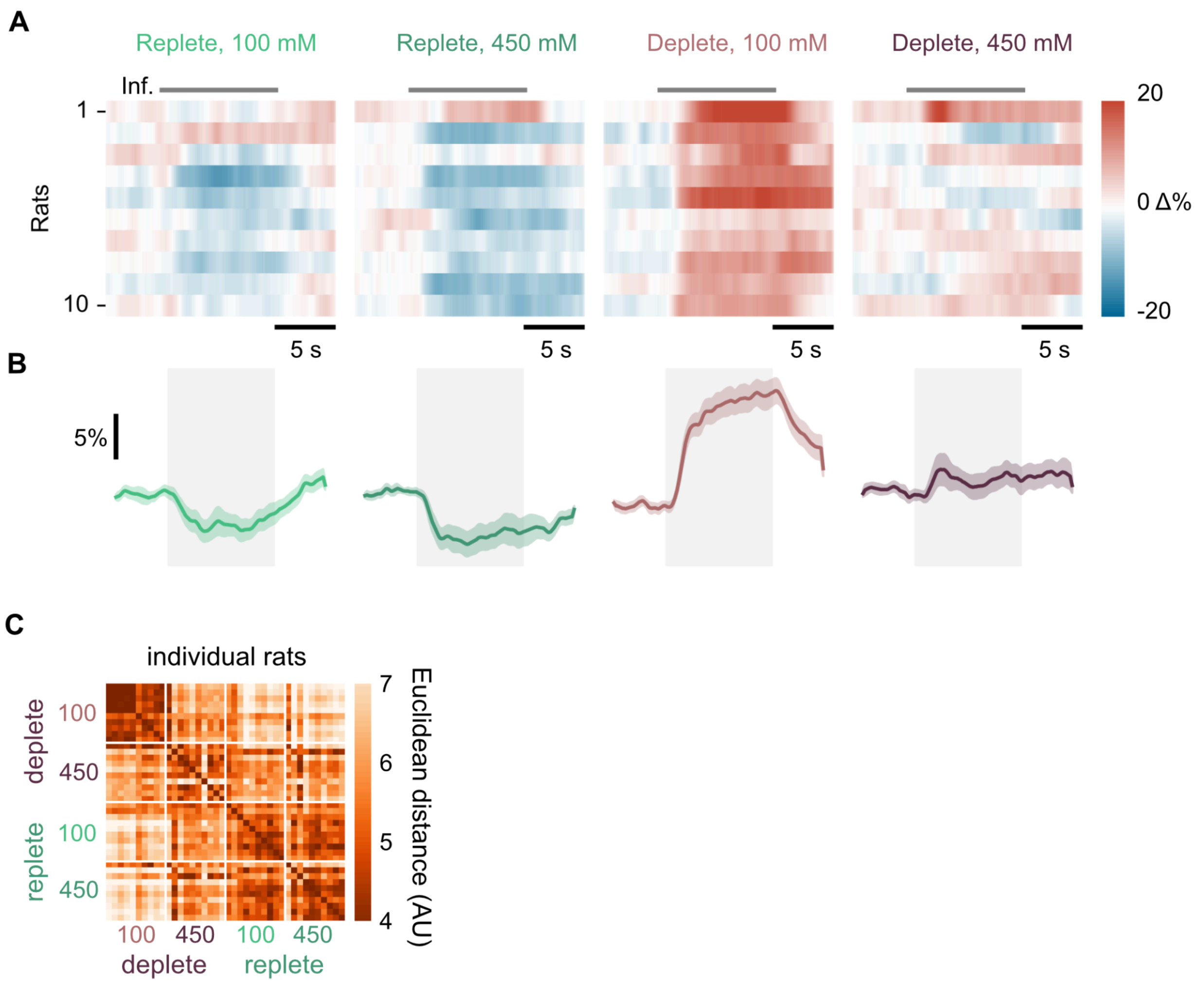


**Figure S2. Rat-by-rat behavioural responses to NaCl in different groups.**

(A) Heatmaps showing probability of appetitive behaviour during intraoral infusions for individual rats. Data are the mean of all trials averaged together. Gray bars show time of infusion.

(B) Plots showing mean +/- SEM of appetitive behaviour probability for all rats. The grey shaded area shows infusion time.

(C) Heatmap showing Euclidean distance of each individual rat's behavioural response across trials, relative to all other rats. Rats are organised by group. Darker colours indicate rats that are closer together.

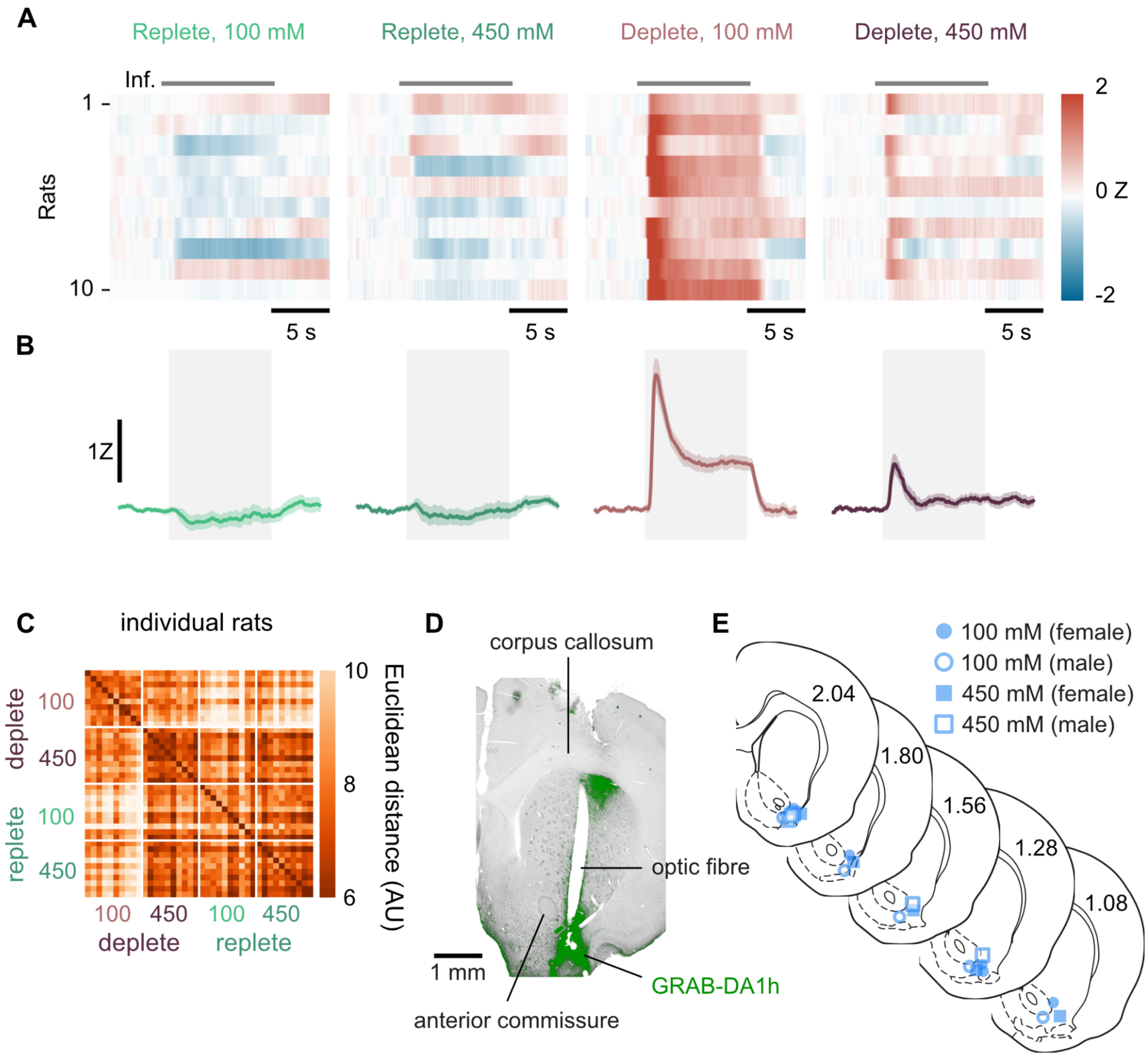


**Figure S3. Rat-by-rat dopamine responses to NaCl in different groups.**

(A) Heatmaps showing dopamine response during intraoral infusions for individual rats. Data are mean of all trials averaged together. Gray bars show time of infusion.

(B) Plots showing mean +/- SEM of dopamine response for all rats. Grey shaded area shows infusion time.

(C) Heatmap showing Euclidean distance of each individual rat's dopamine response across trials, relative to all other rats. Rats are organised by group. Darker colours indicate rats that are closer together.

(D) Representative histology image showing placement of optic fibre and virus expression (shaded green) in nucleus accumbens lateral shell.

(E) Schematic showing optic fibre placements for all rats in study. Numbers on sections are mm from Bregma in anterior-posterior axis.

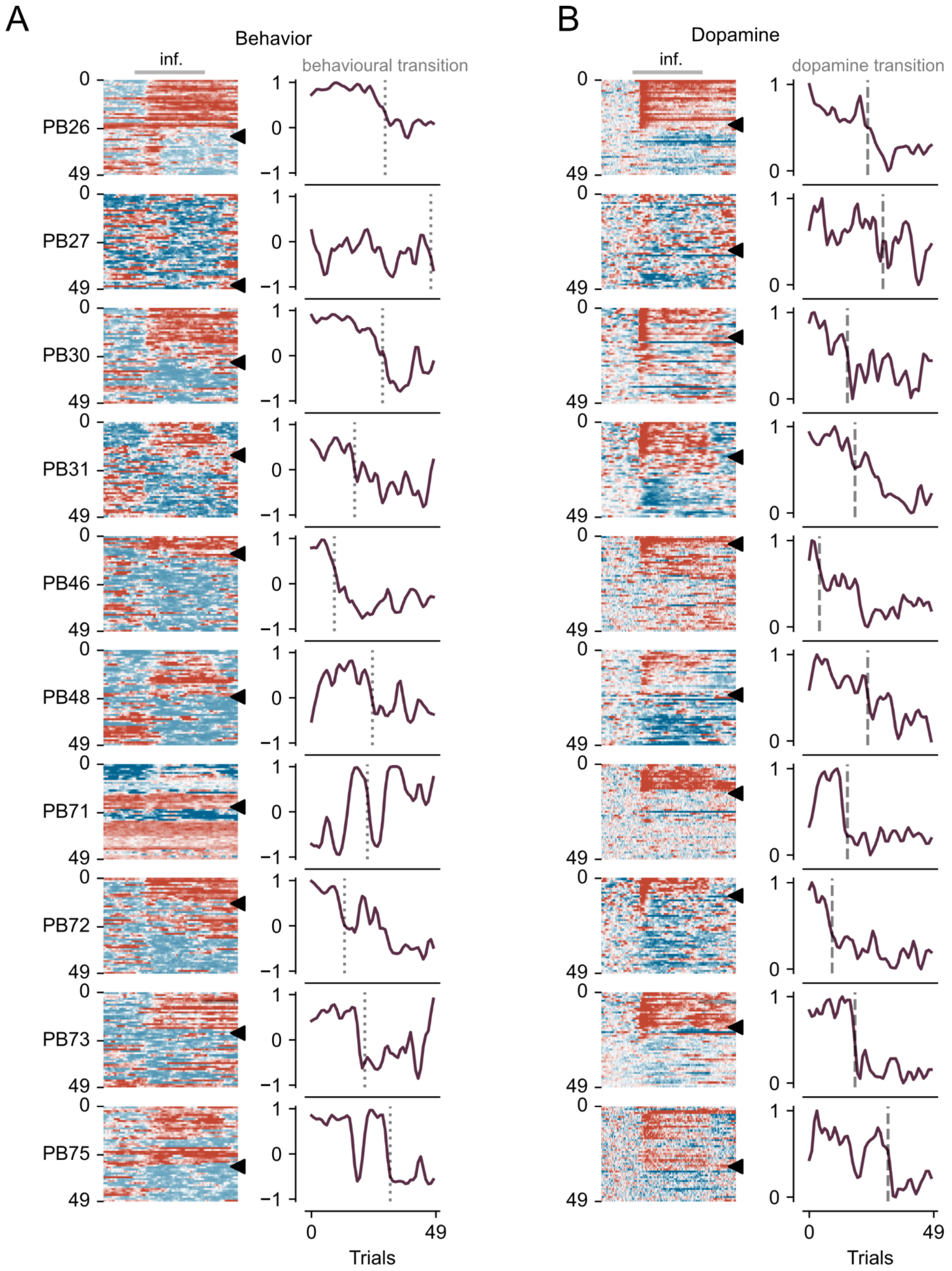


**Figure S4. Behavioural and dopamine responses in individual rats.**

(A) Heatmaps in the left column show appetitive probability across all 49 trials. Line plots in the right column show appetitive probability across trials with the dotted line showing the calculated transition point of the behavioural signal across trials based on maximum velocity of change.

(B) As in A but for dopamine release with dashed line showing the calculated transition point of the dopamine signal across trials based on maximum velocity of change.

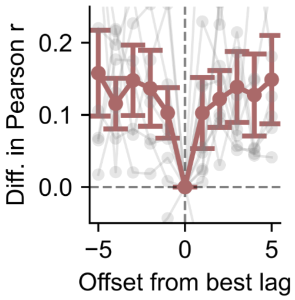


**Figure S5. Sharpness of optimal lag after cross-correlation.**

Plot shows that optimal lag for each rat shows better correlation value (Pearson *r*) than at adjacent lag values. Thick line is mean and grey lines show change for individual rats.

**Table S1 - Fits of trial-by-trial data for appetitive probability**

| Group | Best type of fit | *r* | *p* |
|---|---|---|---|
| Replete - 0.1 M | Linear | 0.484 | <0.001 |
| Replete - 0.45 M | Linear | -0.092 | 0.528 |
| Deplete - 0.1 M | Linear | -0.204 | 0.160 |
| Deplete - 0.45 M | Sigmoidal | 0.902 | <0.001 |

**Table S2 - Fits of trial-by-trial data for dopamine AUC**

| Group | Best type of fit | *r* | *p* |
|---|---|---|---|
| Replete - 0.1 M | Linear | -0.173 | 0.236 |
| Replete - 0.45 M | Sigmoidal | 0.737 | <0.001 |
| Deplete - 0.1 M | Linear | -0.840 | <0.001 |
| Deplete - 0.45 M | Sigmoidal | 0.958 | <0.001 |
